# NanoReviser: An Error-correction Tool for Nanopore Sequencing Based on a Deep Learning Algorithm

**DOI:** 10.1101/2020.07.25.220855

**Authors:** Luotong Wang, Li Qu, Longshu Yang, Yiying Wang, Huaiqiu Zhu

## Abstract

Nanopore sequencing is regarded as one of the most promising third-generation sequencing (TGS) technologies. Since 2014, Oxford Nanopore Technologies (ONT) has developed a series of devices based on nanopore sequencing to produce very long reads, with an expected impact on genomics. However, the nanopore sequencing reads are susceptible to a fairly high error rate owing to the difficulty in identifying the DNA bases from the complex electrical signals. Although several basecalling tools have been developed for nanopore sequencing over the past years, it is still challenging to correct the sequences after applying the basecalling procedure. In this study, we developed an open-source DNA basecalling reviser, NanoReviser, based on a deep learning algorithm to correct the basecalling errors introduced by current basecallers provided by default. In our module, we re-segmented the raw electrical signals based on the basecalled sequences provided by the default basecallers. By employing convolution neural networks (CNNs) and bidirectional long short-term memory (Bi-LSTM) networks, we took advantage of the information from the raw electrical signals and the basecalled sequences from the basecallers. Our results showed NanoReviser, as a post-basecalling reviser, significantly improving the basecalling quality. After being trained on standard ONT sequencing reads from public *E. coli* and human NA12878 datasets, NanoReviser reduced the sequencing error rate by over 5% for both the *E. coli* dataset and the human dataset. The performance of NanoReviser was found to be better than those of all current basecalling tools. Furthermore, we analyzed the modified bases of the *E. coli* dataset and added the methylation information to train our module. With the methylation annotation, NanoReviser reduced the error rate by 7% for the *E. coli* dataset and specifically reduced the error rate by over 10% for the regions of the sequence rich in methylated bases. To the best of our knowledge, NanoReviser is the first post-processing tool after basecalling to accurately correct the nanopore sequences without the time-consuming procedure of building the consensus sequence. The NanoReviser package is freely available at https://github.com/pkubioinformatics/NanoReviser.

## 1 Introduction

Oxford Nanopore Technologies (ONT) has recently shown the ability to use nanopores to achieve long-read sequencing, which has a tremendous impact on genomics (Brown and Clarke, 2016). The small-sized MinION device introduced by ONT is able to translate long sequencing reads from electrical signals produced by a single-stranded molecule of DNA passing through a nanopore (Lu et al., 2016). MinION is much cheaper than the other available sequencers and allows for sequencing to be carried out in real time, and hence has great prospects for use in many areas of genomics analysis such as variation detection and genome assembly (Leggett and Clark, 2017; Ameur et al., 2018; Pollard et al., 2018).

Although it is possible to produce long sequencing reads when using the MinION device with large arrays of nanopores, nanopore sequencing reads do show a fairly high error rate which could be introduced by abnormal fluctuations of the current or by the improper translation from raw electrical signals into DNA sequence, which has been sought to be corrected by the basecalling procedure developed by ONT (Rang et al., 2018). Several basecallers have been proposed since MinION was launched. Albacore (Loman et al., 2015), Guppy and Scrappie are basecallers provided by ONT. Notably, both Albacore and Guppy are only available to ONT’s commercial customers, while Albacore is the default basecaller. Scrappie, a technology demonstrator provided by ONT, actually combined with two basecallers in one: scrappie events and scrappie raw, which can only provide fasta output and was not trainable. Nanocall (David et al., 2017) predicted the DNA sequences using hidden Markov models (HMMs), but could not detect long homopolymer repeats. DeepNano (Boža et al., 2017), the first basecaller based on deep learning and one applying recurrent neural networks (RNNs) to predict DNA bases, showed that a deep learning model could make a crucial contribution to the basecalling performance. However, DeepNano was developed mainly based on the datasets sequenced using R7.3 and R9.0 flow cells, which are no longer used. Later on, BasecRAWller (Stoiber et al., 2017b), designed to use the raw outputs of nanopores with unidirectional RNNs to basecall DNA bases, was shown to be able to process the electrical signals in a streaming fashion but at the expense of basecalling accuracy. Chiron (Teng et al., 2018), designed to adopt a combination of convolution neural network (CNN) and RNN methodologies together with a connectionist temporal classification decoder (CTC decoder) that was successfully used in machine translation, achieved a high basecalling accuracy. Despite this high accuracy, Chiron, however, required a high much computer time to get these accurate results. Notably, all of these tools were designed to focus on basecalling but not sequencing error correction. The only tool proposed previously for MinION sequencing error correction, nanoCORR (James Gurtowski, 2016), requires additional next-generation sequencing short read sequences, and thus is not a practical tool for error correction in its true sense.

In this study, we developed NanoReviser, the first error-correction tool for a MinION nanopore sequencing platform. To correct errors introduced by the default basecallers provided by the manufacturer, our tool was designed to first take both the original electrical signals and base sequences as the inputs. In the module designed for the tool, we used a CNN (Kalchbrenner et al., 2014) to extract the local patterns of the raw signals, and a highly powerful RNN (Pascanu et al., 2013) and bidirectional long short-term memory networks (Bi-LSTMs) (Hochreiter and Schmidhuber, 1997; Schuster and Paliwal, 1997) to determine the long-term dependence of the bidirectional variation of the raw electronical signals on DNA strand passing through the nanopore hidden in the basecalled sequences. Although the module was trained on a small dataset from *Escherichia coli* K12 MG1655 (Jain et al., 2017) and human NA12878 (Jain et al., 2018) sequences, it was shown to be able to correct errors of sequencing for many other genomes as long as corresponding training sets were provided. Furthermore, since studies have shown that DNA modification information could be detected by analyzing the electrical signals and since both the DNA sequences and the DNA modifications could be determined at the same time using the Oxford Nanopore sequencing (Stoiber et al., 2017a), we analyzed the basecalling results for the methylated bases and trained our module with the modified data. We believe that NanoReviser will find significant use in genome analysis when ONT sequencing becomes, as expected, widely used in the future.

## 2 Materials and Methods

### 2.1 Data sets

We performed this study with data from two public datasets: human data from the Nanopore WGS Consortium (Stoiber et al., 2017a) and *E. coli* data from the MinION Analysis and Reference Consortium (MARC) (Jain et al., 2017). The Nanopore WGS Consortium sequenced the genome of the human sample CEPH1463 (NA12878/GM12878, Ceph/Utah pedigree) using 1D ligation kits with R9.4 chemistry. The sequencing of this data was basecalled using Albacore release v2.0.2. We trained our module on the chromosome 20 data and validated our module on the rel3 NA12878 genome data. MARC provided the whole genome sequencing of *E. coli* K12 MG1655 using 1D ligation kits and Rapid kits with R9.0 chemistry, and the basecaller used for these samples was Albacore release v0.8.4. A summary of the two datasets is listed in Table 1.

**Table 1.**
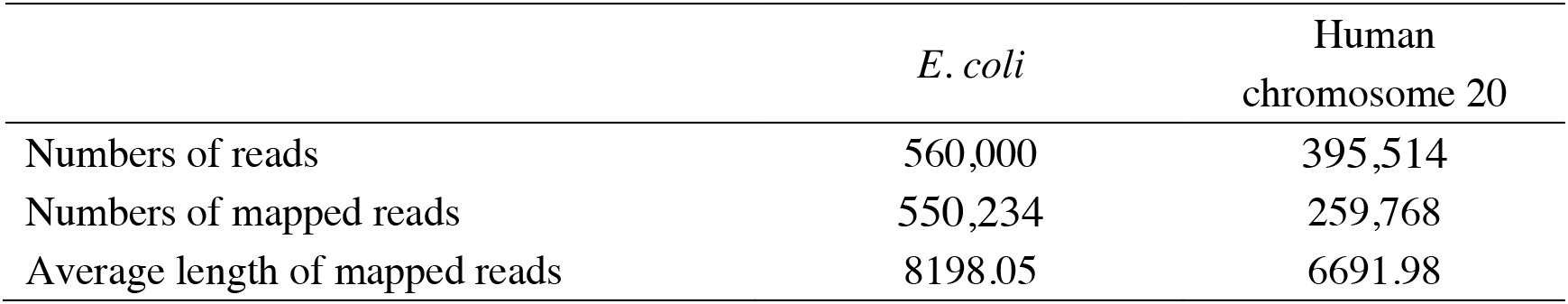
Summary of the experimental datasets for training NanoReviser

The sequence of the *Escherichia coli* K12 MG1655 genome may be accessed at the NCBI under Refseq accession number NC_000913.3 and the sequence of the *Homo sapiens* NA12878 genome was downloaded from the NCBI GenBank assembly accession GCA_002077035.3. The annotated file of the modifications in the *E. coli* genome was downloaded from the MethSMRT database (Ye et al., 2017).

### 2.2 Preprocessing

#### 2.2.1 Re-segmentation of events during sequencing

The raw signals generated by the process of DNA passing through the nanopores were basecalled into DNA sequences immediately by default. The raw signals, the files used during the basecalling procedure, and the output file in fastq format were packed together into a fast5 file as the final output of the sequencing. Based on the files used during the basecalling procedure, which were the event files, we re-segmented the events provided by the default Albacore basecaller.

A period of raw signals that can be translated into a particular length of DNA was defined when using ONT procedures as an event, and a series of parameters were used to describe this event (David et al., 2017). Although the parameters in an event defined by the different basecallers differed, the common parameters were *mean, start, stdv, length, model_state, move, weight,* and *p_model_state* — with *model_state* referring to the corresponding predicted DNA sequence of the event, *start* being the starting time point of the event, *length* being the length of the raw signals corresponding to the event, *mean* and *stdv* being the basic statistics of this period of raw signals, *move* describing the movement of the DNA in nanopores and having only three values (0 standing for the DNA strand staying in the nanopore and 1 and 2 standing for the DNA strand moving, respectively, one base or two bases in the nanopore), *weight* and *p_model_state* being essential parameters for basecalling. Based on *move*, we re-segmented the events and calculated the *mean* and *stdv* as the input of our module. The re-segmentation process was crucial for basecalling in the following process.

#### 2.2.2 Labeling and training

We aligned the re-segmented reads against the *E. coli* K12 MG1655 reference genome using GraphMap (Sović et al., 2016), which was designed specifically for the alignment of the nanopore reads, with the default parameters and we labeled the sequencing errors and basecalling errors based on the alignment results. As for the methylation labeling, we aligned the re-segmented reads against the *E. coli* K12 MG1655 reference genome to get the positions of events. Then, we labeled the methylated bases according to the annotation from the MethSMRT database (Ye et al., 2017), which defined three types of methylated areas: m6A stands for N6-methyladenosine, m4C stands for N4-methylcytosine and modified bases.

Notably, the two dataset projects (Nanopore WGS Consortium and MARC) were quite different in many of their sequencing procedures such as library preparation (Rapid kit vs. 1D ligation kit), flow cell version (R9.4 chemistry vs. R9.0 chemistry) and basecaller tools (Albacore v2.0.2 vs. Albacore v0.8.4). Thus, we trained NanoReviser on the *E. coli* data and human data individually. In order to build the training set and validation set, we first divided the reference genome into 5,000 bp intervals. Then we aligned all of the reads and separated them into different intervals based on the alignment results. For the *E. coli* training process, we selected two reads from every interval, which was 2x depth data, to build up the low-coverage training set and we selected the reads located on the genome length of 2,225,000 bp to 2,300,000 bp to build up the local training set: There were 1,854 and 2,412 reads in, respectively, the low-coverage and local *E. coli* training sets.

We used 1x depth reads, which means the reads contained in our selection could cover the genome one times, and the reads located on 28,750,000 bp to 43,750,000 bp of the NA12878 chromosome 20 to build, respectively, the low-coverage training data and local training data for the training process using the human data. The low-coverage training data contained 4,820 reads and the local data contained 4,593 reads.

We used the Adam (adaptive moment estimation) algorithm with the default parameters (Kingma and Ba, 2014) in the training process to perform optimization. NanoReviser iterated through 50 epochs with the low-coverage data, the local data, and the combination of the low-coverage data and local data, which we denoted as NanoReviser (low coverage), NanoReviser (local) and NanoReviser respectively. We trained our models on a high-performance computer (Dell R930, 2019) with four CPU processors (2.1 GHz Intel Xeon E7-4809 v4), four GPU processors (Nvidia Tesla T4) and 18 DDR4 memories (18×32G, 2133MHz). Additionally, we applied multiprocessing implementation in the training procedure of NanoReviser to reduce training time, and the corresponding training time is presented in Table 2.

**Table 2.**
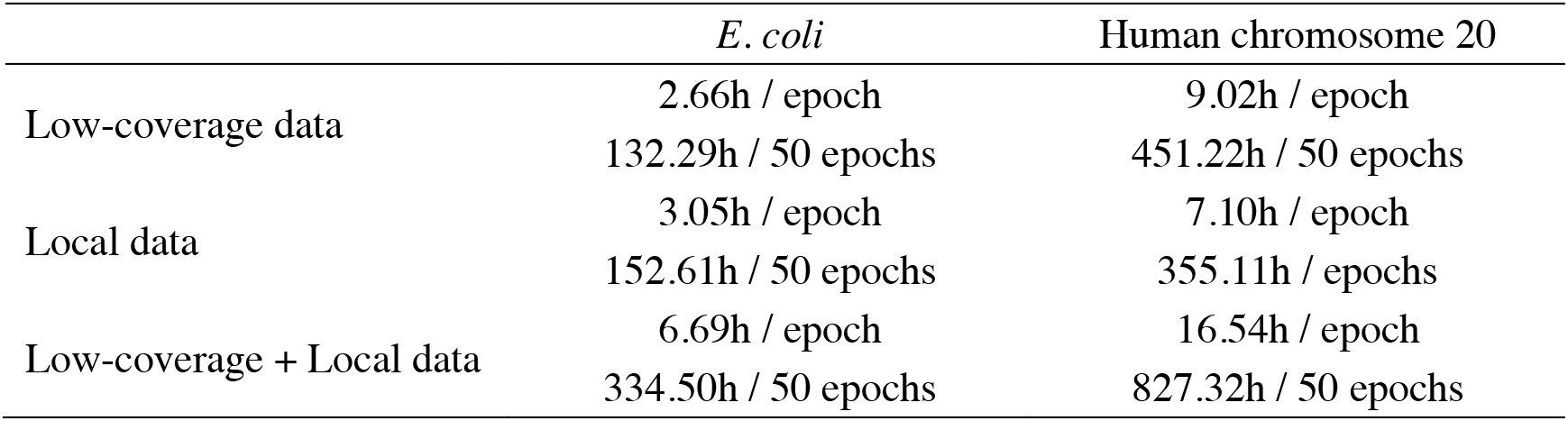
Summary of the training time

### 2.3 Model architecture

As shown in Figure 1A, NanoReviser was designed to be composed of two main models, named model 1 and model 2, respectively, with the main task for model 1 being the learning of the error types from the combination of the reads and electrical signal input and the main goal of model 2 being identification of the bases. Model 1 and model 2 were designed to share a similar model structure, called the main model, which is shown in Figure 1B.

**Figure 1.**
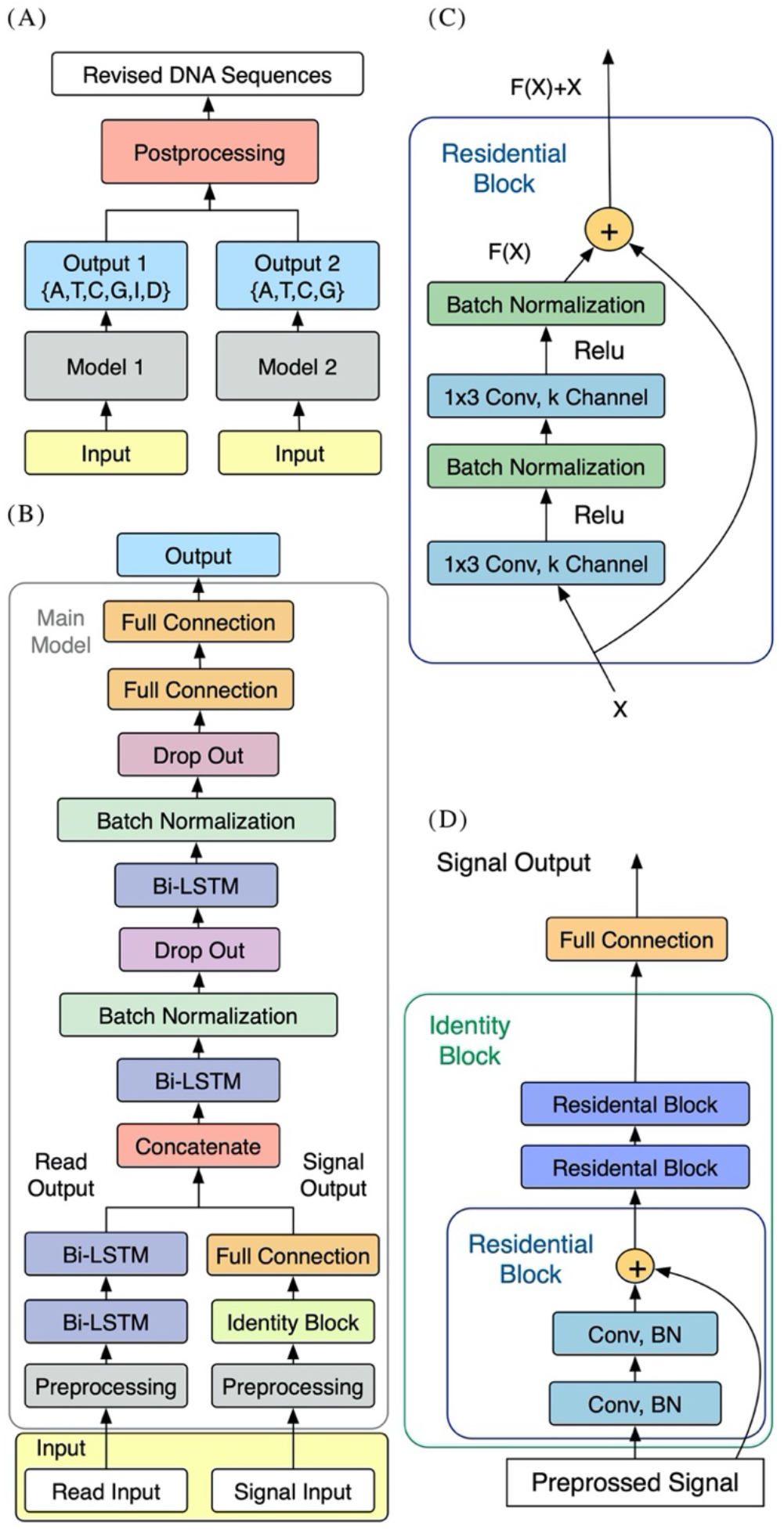
Schematics of NanoReviser. **(A)** Structure of NanoReviser model building. **(B)** Structure of the main model. The preprocessed raw electrical signals (on the right) were passed through Identity Block and joined with the results of the preprocessed read input (on the left), passing through two bidirectional long short-term memory (Bi-LSTM) layers, and then the combination of the raw electrical signal features and read features were fed into the following Bi-LSTM layers. Finally, after the formation of two fully connected layers, the model gave a probability distribution of called bases. **(C)** Structure of Residential Block. Residential Block consisted of two convolutional layers and two batch normalization layers, which were used to accelerate the training speed. Conv stands for a convolutional layer and 1×3 was the size of the kernel used by the convolutional layer. **(D)** Structure of Identity Block. Identity Block consisted of three Residential Blocks. Conv stands for a convolutional layer and BN is the abbreviation for Batch Normalization.

#### 2.3.1 Main model architecture

A simple multi-layer RNN cannot precisely capture the fine-scale changes of the electrical signals caused by DNA polymers passing through nanopores. In order to increase the model capacity, we designed the main model to be similar to a speech recognition deep learning model (Rosenberg et al., 2017) that used multiple CNN layers (Kalchbrenner et al., 2014) and RNN layers (Pascanu et al., 2013). In order to make our module perform well with extremely imbalanced datasets, we also exploited the center loss function (Wen et al., 2016) in our module. Figure 1B shows the architecture of our sequencing error correction system.

In our definition, the event of moment *t* defined as *e*_*t*_ was labeled as *y*_*t*1_ ∈ {A, T, C, G, I, D} in model 1 (with I and D representing, respectively, an insertion and deletion in the CIGAR record of the SAM format file) and labeled as *y*_*t*2_ ∈ {A, T, C, G} in model 2. As for the event of moment *t*, we defined the first input, i.e., the raw electrical signal, as ***s***^(***t***)^ — with ***s***^(***t***)^ corresponding to a time-series of length ***T***^(*t*)^, and with every time slice represented as a vector of normalized raw signal features in the form 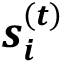 , ***i*** = 0, … , ***T***^(*t*)−1^. We defined the second input as ***x***^(***t***)^ . which consisted of the basecalled bases as well as the mean, standard deviation and length of the event of moment *t*. Besides, the second inputs we used for NanoReviser methylations contained the features mentioned above, as well as the methylated information of the events. These two inputs composed the input as shown in Figure 1B.

To be specific, for model 1 or model 2, first ***s***^(***t***)^ was passed through the multi-layer CNN module and the second input ***x***^(***t***)^ was passed through the unique powerful RNN layers, which are the Bi-LSTM (Hochreiter and Schmidhuber, 1997) layers. Then, the two inputs were joined together as the input of the following multi-layer Bi-LSTM module. Finally, the output ***o***_***t***_ at the *t* moment gave a probability distribution of called bases. Our module was designed with the core idea to convert the input ***s***^(***t***)^ and the input ***x***^(***t***)^ into a final transcription ***y***_***t*1**_ or ***y***_***t*2**_. Notably, we used center loss to help supervise the learning of our module.

#### 2.3.2 Convolutional neural network layers

Since our electrical signal input was a one-dimensional time-series sequence, we used a one-dimensional CNN layer with a filter size and stride set of 3 and 1, respectively. Then we applied the rectified linear unit (ReLU) function (Zhao et al., 2017) on the outputs of the CNN layer, which was the most common activation operation in the CNN models (Table 3). We also used the batch normalization to accelerate the training process.

**Table 3.**
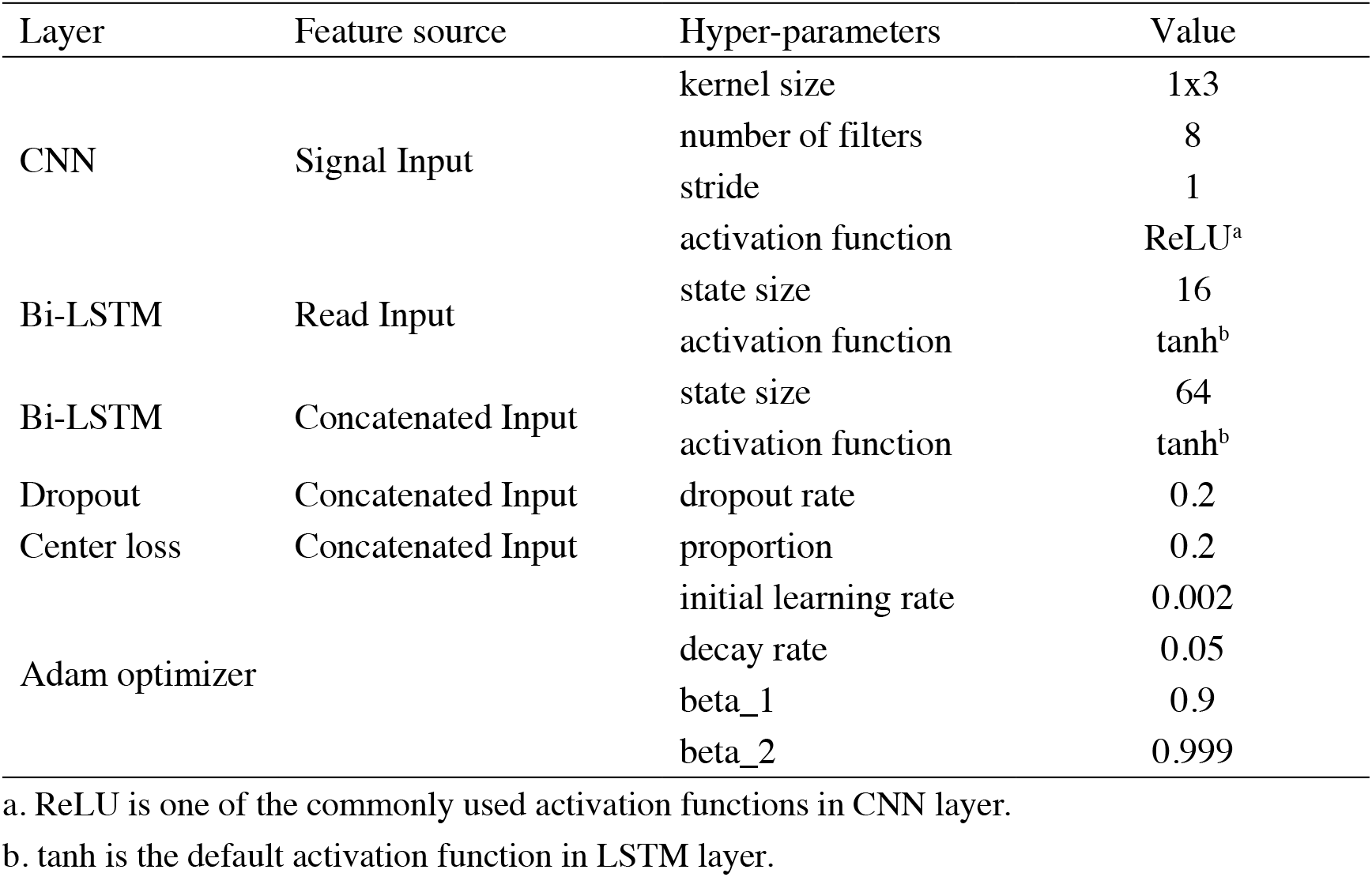
Hyper-parameters of NanoReviser

CNN is good at learning multi-level representations, but the traditional deep CNN faces a degradation problem (LeCun et al., 2015). Thus, in the current study, we used the residual CNN (He et al., 2015) to extract features from the raw signals. The residual network contained direct links between the higher layer outputs and the lower layer inputs, and the residual block in our module is illustrated in Figure 1C.

We used three residential blocks to build an identity block (Figure 1D) just like the ResNet model (He et al., 2015). Indeed, the first input ***s***^(***t***)^ went through the identity block and then was mixed with the second handled inputs.

#### 2.3.3 Recurrent neural network layers

Compared with the traditional recurrent layers, the LSTM model (Hochreiter and Schmidhuber, 1997) has been shown to be more effective, especially for the task of learning long-term dependencies (Yue and Wang, 2018). We applied the tanh function (Yue and Wang, 2018)on the outputs of the LSTM layer, which was the default activation operation in the LSTM models (Table 3). Since DNA polymers near the nanopores could contribute to changes in the electrical signal, a Bi-LSTM model (Schuster and Paliwal, 1997) allowed us to look at the bases both far ahead and far behind at will. Thus, we recruited the Bi-LSTM model in our module. Specifically, the second input, ***x***^(***t***)^, was fed into two Bi-LSTM layers, combined with batch normalization. Then, the latter Bi-LSTM layer received the combination of the handled signal data and read data. In order to avoid overfitting, we used dropout layers (Bouthillier et al., 2015). After formation of the two fully connected layers, model 1 finally output the likelihood of the input event as A, T, C, G, I or D and model 2 output the likelihood of the input event as A, T, C or G.

#### 2.3.4 Loss function

In order to make our module perform well with extremely unbalanced data, we recruited the center loss function (Wen et al., 2016) into the commonly used softmax cross-entropy loss function. According to Wen, the softmax cross-entropy loss function merely considers the inter-class distance, whereas the main idea of the center loss function is to reduce the intra-class distance during the training process.

The softmax cross-entropy loss function is normally defined as

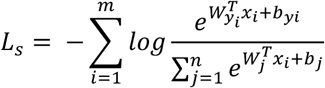

where *x*_*i*_, which belongs to the *y*_*i*_th class, is the ***i*** th feature of the mini-batch inputs with *m* samples. Furthermore, *W and b* are the weights and bias term in the last fully connected layer, and *n* is the number of classes. Obviously, the model could learn to distinguish the samples from different classes by minimalizing the softmax cross-entropy loss.

The center loss function used in our current work was expressed using the equation

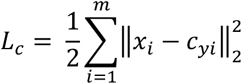

where *m* is the mini-batch number and *c*_*yi*_ denotes as the center of features. The loss function we used in our module was just like the one Wen (Wen et al., 2016) used in his work and was represented as:

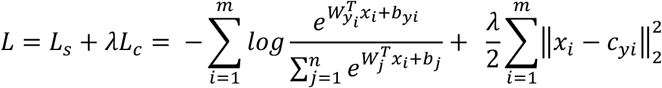

Here, *λ* is a hyper parameter used to control the proportion of the softmax cross-entropy loss and the center loss. We set *λ* to be 0.2.

### 2.4 Evaluation criterion

Due to the sequencing protocols and default basecallers being quite different between the *E. coli* (K12 MG1655) and human (NA12878) datasets, we evaluated NanoReviser on both the *E. coli* and human datasets separately. We randomly used 100 reads that were not contained in training set from *E. coli* data apart from the training set to evaluate our module. As for the NA12878 datasets, 100 reads excluded from the training set from human NA12878 chromosome 20 and 2,100 reads from the other chromosomes (100 reads from each chromosome apart from chromosome 22, which is not provided in the rel3 data of the Nanopore WGS Consortium) were chosen to evaluate the performance of our module, and compared this performance to those of other modules.

The evaluation criteria we used were the deletion rate, the insertion rate, the mismatch rate and the error rate, where

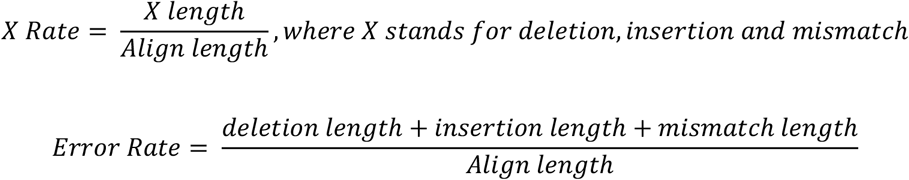

and

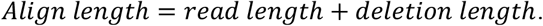

According to methylation annotation of the *E. coli* genome on the MethSMRT database(Ye et al., 2017), there are three types of methylated areas: m6A, m4C and modified bases. Therefore, the deletion rate, the insertion rate, and the mismatch rate on the kth type of methylated areas could be denoted as:

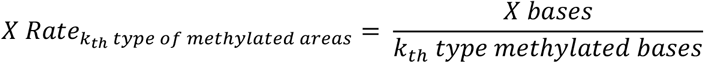

Where *X* stands for deletion, insertion and mismatch. The error rate on the k type methylated areas could be denoted as:

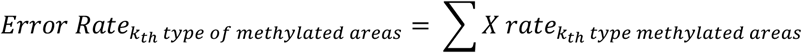

Thus, the total deletion rate, the insertion rate, the mismatch rate and the error rate on the methylated areas could be denoted as:

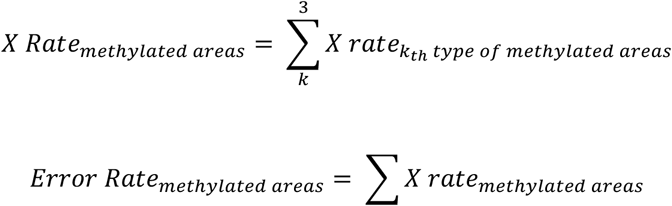

Where *X* stands for deletion, insertion and mismatch.

We compared NanoReviser with DeepNano (Boža et al., 2017), Chiron Versions 0.6.1.1 (Teng et al., 2018) and Scrappie 1.4.0 (https://github.com/nanoporetech/scrappie)

## 3 Results and discussion

### 3.1 Implementation of NanoReviser

NanoReviser used the two outputs from two deep learning models to compute the final DNA sequences as shown in Figure 1A. Specifically, when model 1 or model 2 predicted the same base as the original read base, the original read base was preserved; whereas when model 1 and model 2 each predicted the same base as each other but different than the original read base, the read base was corrected. NanoReviser directly deleted the read base when model 1 considered the read base to be an insertion and model 2 predicted the same base as the original read base. When NanoReviser predicted a base as a deletion, the base inferred by model 2 was added to the DNA sequence. Except for these situations, NanoReviser tended to preserve the original read bases to avoid introducing new errors.

Since model 1 and model 2 share a similar structure, we have tested the parameters on the model 1 and used the same parameters for model 2. Table 3 summarizes the chosen hyper-parameters for NanoReviser. The batch size is set to 256 and the training sets were shuffled at the beginning of every training round. As described in the previous studies, the presence of multiple bases in the nanopore could contribute to the raw electrical signal produced and used to predict the identity of the central base(Rang et al., 2018). Although LSTM could relatively well memorize the long-term relationship between the surrounding bases and the central base in the nanopore, processing the whole nanopore sequencing read is still beyond the ability of LSTM. Therefore, the inputs must be divided into small pieces, and we tested window sizes ranging from 9 to 15. We observed that the model 1 performed the best when the window size was 13 and we inferred from our results that the model converged after 50 rounds of training (Figure 2). Since window size, training epochs and the data size of the training data are crucial for the final training result, these parameters could be set by users when training NanoReviser.

**Figure 2.**
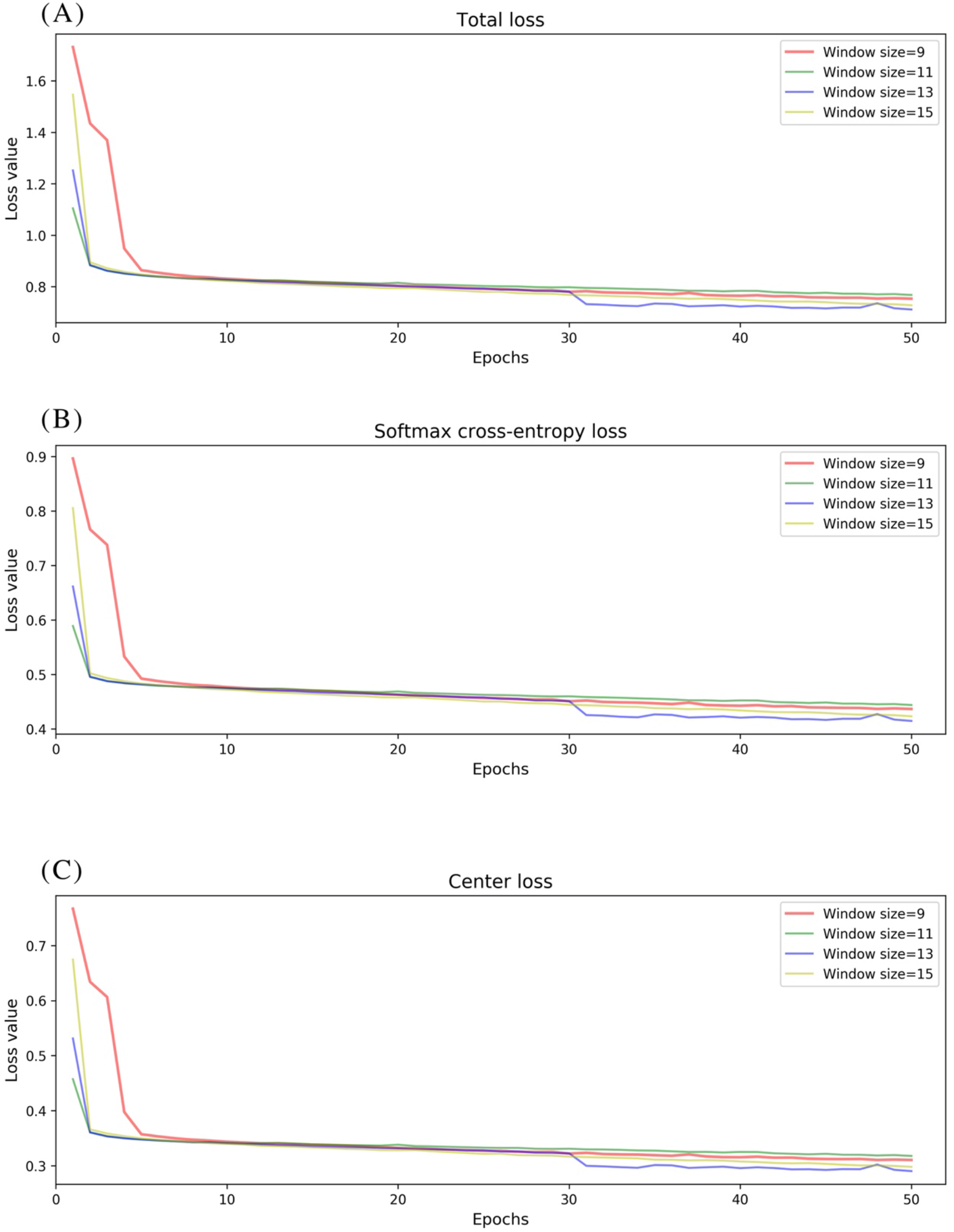
NanoReviser fitting performances for various window sizes and over many iterations. **(A)** Total loss value. **(B)** Softmax cross-entropy loss value. **(C)** Center loss value.

### 3.2 Performance comparison on the *E. coli* dataset

To date, 560,000 nanopore sequencing reads have been collected in the *E. coli* dataset and the sequencing depth of the *E. coli* dataset used was near 650x. We randomly selected 100 reads from the 560,000 reads for evaluation. The details of the NanoReviser performance are presented in Table 4. Specifically, we separated 2x depth reads to train our module and the method was marked as NanoReviser (low coverage). Then, we trained our module including only the reads located in a particular area of the *E. coli* genome and the method was marked as NanoReviser (local). Moreover, we trained our module with a combination of both low-coverage reads and local reads, and this method was marked as NanoReviser. Both NanoReviser (local) and NanoReviser (low coverage) performed better than did the default basecaller, i.e., the Albacore program, as well as the Chiron program, which is currently considered to be the most accurate open-source basecaller (Wick et al., 2019). Unfortunately, although DeepNano could yield all the reads in test data, all the reads could not be aligned to the *E. coli* genome. Thus, DeepNano was excluded from the comparison on the *E. coli* data. When compared with the latest basecaller Scrappie, NanoReviser performed better than both the Scrappie events and Scrappie raw model. Notably, scrappie raw achieved a good performance at the expense of fewer reads production, which generated only 90 reads on the 100 reads test sets. After 50 rounds of training, NanoReviser reduced the error rate by over 5% compared with the default basecaller Albacore and achieved the best performance on the test set.

**Table 4.**
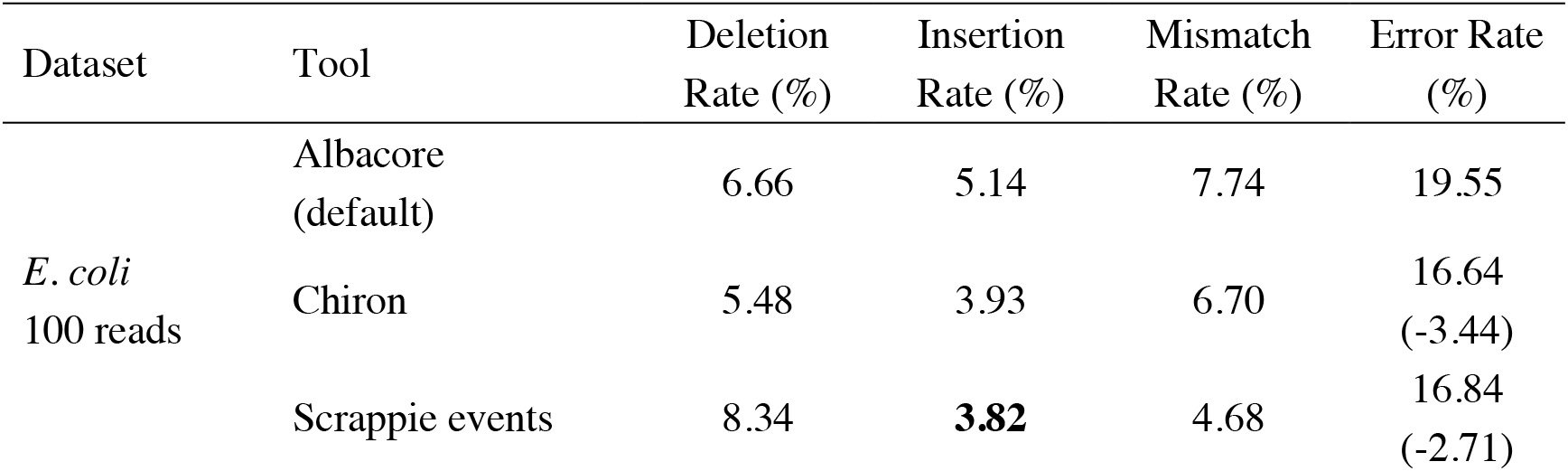
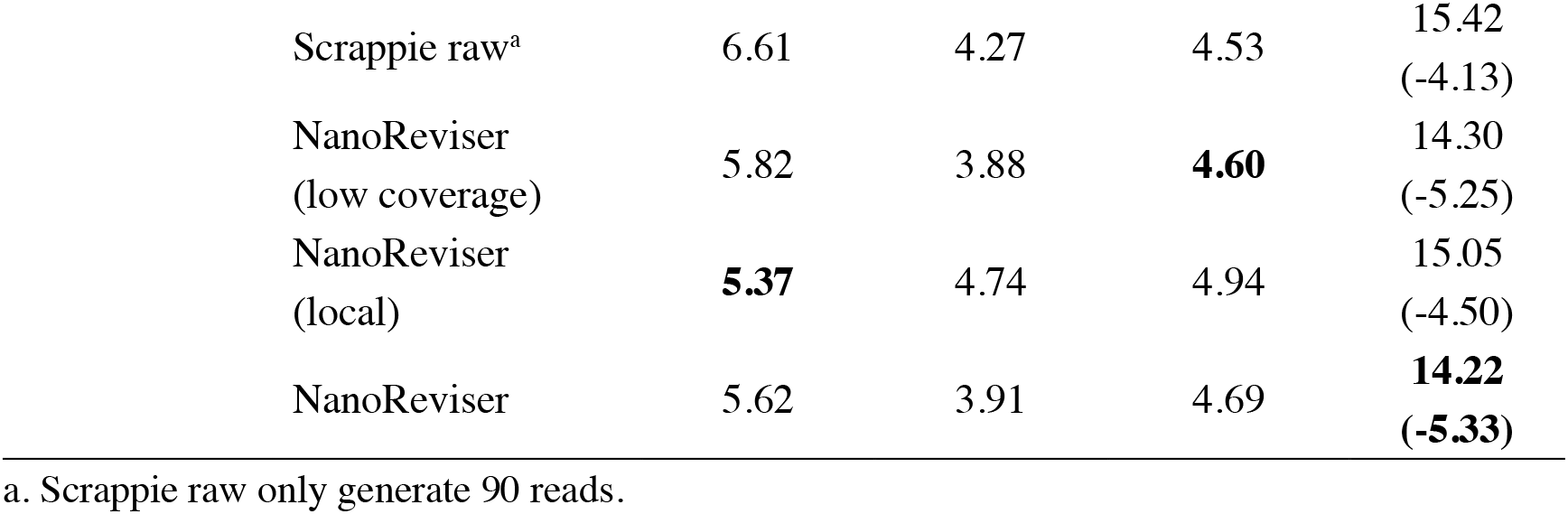
Performances of various tools on *E. coli* genome sequencing data

| Dataset | Tool | Deletion Rate (%) | Insertion Rate (%) | Mismatch Rate (%) | Error Rate (%) |
| --- | --- | --- | --- | --- | --- |
| *E. coli*<br>100 reads | Albacore (default) | 6.66 | 5.14 | 7.74 | 19.55 |
|  | Chiron | 5.48 | 3.93 | 6.70 | 16.64<br>(-3.44) |
|  | Scrappie events | 8.34 | **3.82** | 4.68 | 16.84<br>(-2.71) |

|  |  |  |  |  |
| --- | --- | --- | --- | --- |
| Scrappie raw<sup>a</sup> | 6.61 | 4.27 | 4.53 | 15.42<br>(-4.13) |
| NanoReviser<br>(low coverage) | 5.82 | 3.88 | **4.60** | 14.30<br>(-5.25) |
| NanoReviser<br>(local) | **5.37** | 4.74 | 4.94 | 15.05<br>(-4.50) |
| NanoReviser | 5.62 | 3.91 | 4.69 | **14.22**<br>**(-5.33)** |

a. Scrappie raw only generate 90 reads.

Notably, we found that although NanoReviser improved the basecalling accuracy in general regardless of the data on which it was trained, the training data chosen nevertheless did slightly influence the type of error identified by the model. It seemed that NanoReviser best recognized the patterns of insertion errors from the low-coverage training sets whereas it was most sensitive to the deletion error type when trained on the local reads. Theoretically, the insertion error and the deletion error may be thought of as “two sides of a coin”, i.e., with an expectation that a model tending to be more sensitive to the deletion error type would introduce more insertion errors, and similarly a model tending to be more sensitive to the insertion error type would introduce more deletion errors. When trained with the combined datasets including both low-coverage data and local data, NanoReviser generally achieved a balanced sensitivity to the insertion and deletion error types.

### 3.3 Performance comparison on the human dataset

Compared with the *E. coli* dataset, more advanced sequencing applications were involved in the sequencing of the NA12878 dataset. A total of over 13 million reads have been generated from several laboratories and the sequencing depth of the human dataset used was about 28x. We used the reads from chromosome 20 to train the NanoReviser. In order to test the generalization ability of the NanoReviser, we randomly selected 2,200 reads from chromosome 1-20 and chromosome X of the NA12878 dataset for evaluation. The results of these evaluations are reported in Table 5.

**Table 5.**
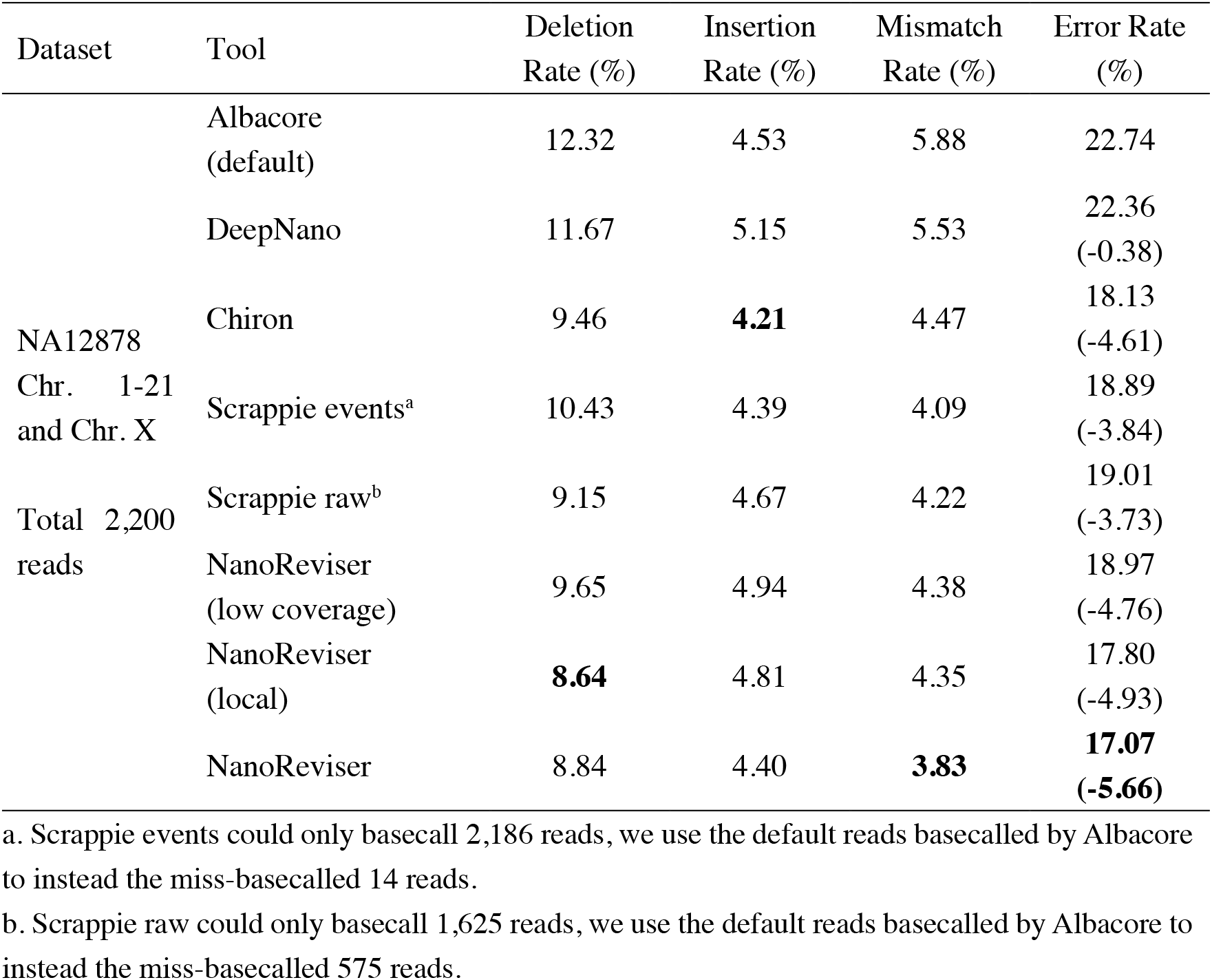
Performances of various tools on human NA12878 genome sequencing data

We used 1x depth reads to train the model and the method was marked as NanoReviser (low coverage) in Table 5. Then, we trained our module including only the reads located on a particular area and the method was marked as NanoReviser (local) in these tables. In both the low-coverage and local training methods, NanoReviser provided better performances than did the default basecaller Albacore. Since both Scrappie events and Scrapie raw could not produce all the reads contained in the test data, we used the reads basecalled by the default basecaller Albacore to make sure all the tools could yield the whole reads contained in the test datasets. As it is shown in Table 5, NanoReviser trained with the local and low-coverage data achieved the best performance, with an error rate of 17.07%, compared to the error rate of 22.74% for the default basecaller and error rate of 18.13% for the Chiron program.

When trained on the *E. coli* dataset, NanoReviser showed the same error recognition pattern as it did when trained on the human dataset, i.e., learning more about insertion errors from the low-coverage data and more about deletion errors from the local data. Moreover, unlike the Illumina sequencing technology whose sequencing errors have been shown to be random (Besser et al., 2018), Oxford Nanopore sequencing technology showed more deletion type errors both when trained on the *E. coli* data and human data. The deletion error rates were observed to be higher than the other types of error rates regardless of the kind of basecalling or revision tools used. Also, for all tools tested, we noticed that the deletion error rates for the human dataset were much higher than those for the *E. coli* dataset. Although the deletion error was indicated to be a serious issue for the Oxford Nanopore sequencing, sacrificing the insertion error rate in order to detect more deletion errors, from comparing the performance of NanoReviser (local) with the performance of NanoReviser (low coverage), was concluded to not be warranted. Fortunately, NanoReviser could learn the tradeoff between detecting deletion errors and insertion errors through a combination of using low-coverage data and local data, and in this way increase the overall performance.

### 3.4 Methylation and error revision on the *E. coli* dataset

Studies have shown that methylation could severely affect the basecalling (Feng et al., 2015; Jain et al., 2016; Simpson et al., 2017; Rang et al., 2018). Therefore, we analyzed the error rates of the methylated bases of the *E. coli* genome. The Oxford Nanopore sequencing technology did encounter troubles in this regard. The overall error rate for the *E. coli* dataset was 19.55%, whereas the error rate for the methylated bases was over 34% and the mismatch error rate was as high as 28.53% (see Figure 3 and Table 5). The deletion error rate seemed to be similar to the overall genome deletion error rate whereas no insertion error happened for the methylated bases

**Figure 3.**
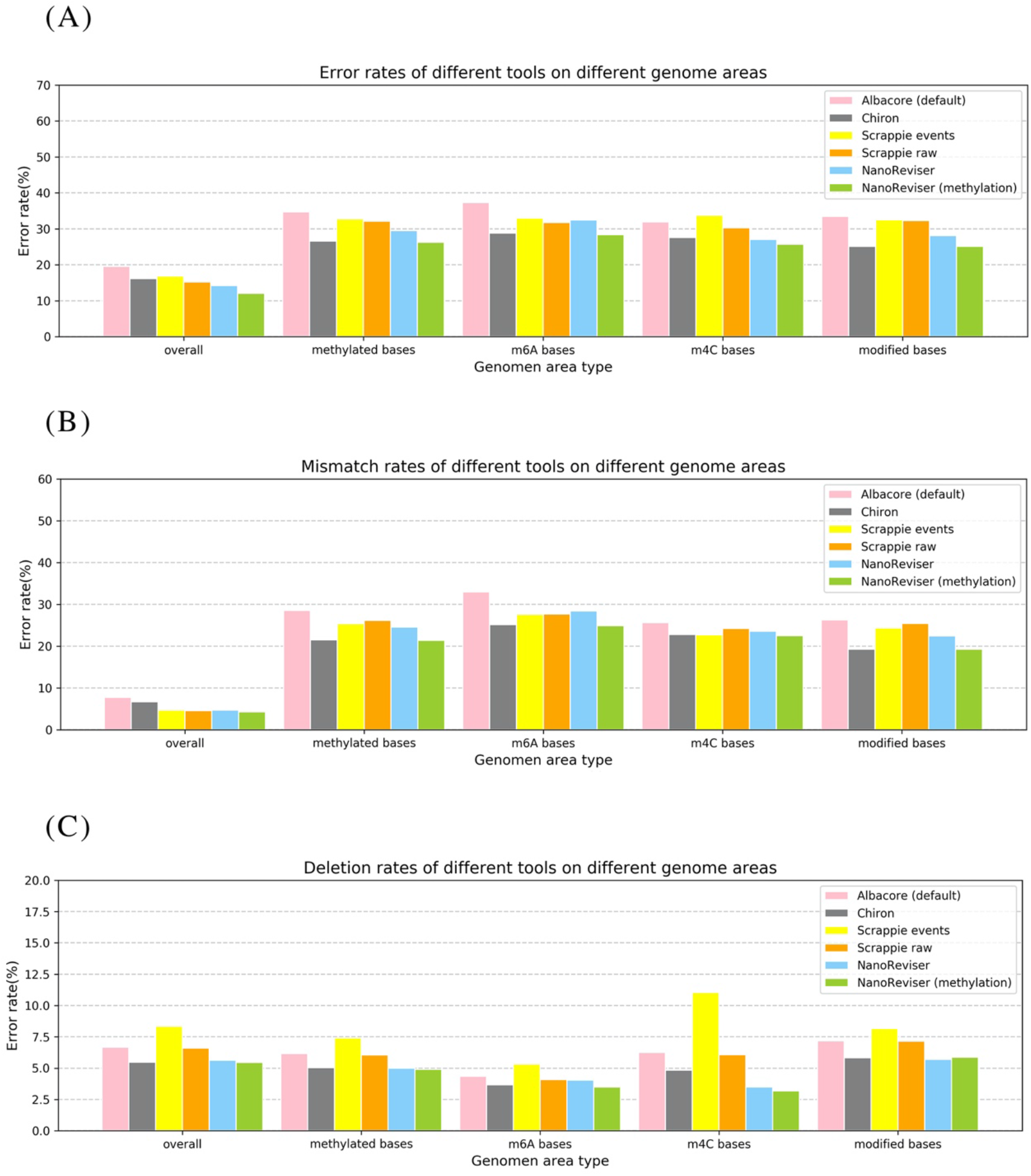
Error rates of different error types on different genome areas. **(A)** Overall error rate. **(B)** Mismatch error rate. **(C)** Deletion error rate. NanoReviser (methylation) stands for NanoReviser trained with methylation information.

Since, promisingly, a tool was able to detect the modified bases at the same time as basecalling, we annotated the methylated areas of the reads based on the MethSMRT database (Ye et al., 2017) and took the methylation information as an additional input feature to train the model. We used the same 100-read *E. coli* test set here as described above, and the results are presented in Table 6. During the NanoReviser (methylation) training process, we used the combination of the low-coverage data and local data we had used in training NanoReviser without methylation. Interestingly, it seems that all tools reduced the error rate by revising the mismatch errors when basecalling the methylated bases. The analysis of insertion rates of different tools on the methylated bases was excluded from Figure 3 and Table 6 because all tools either translate methylated bases into other bases or skip methylated bases. In other words, insertion errors never happened on the methylated bases. As for the methylated area, the overall error rate of NanoReviser without the methylation information was observed to be higher than Chiron, yet was also observed to be lower than the overall error rates of Scrappie raw and Scrapie event (Figure 3). However, by taking the methylation positions into consideration, NanoReviser (methylation) reduced the error rate by about 7.5% compared with the default basecalling result and decreased the error rate by over 2% compared with the model without the methylation information, which achieved the best performance on all kinds of methylated bases including the m6A bases, m4C bases, and the modified bases.

**Table 6.** Performances of various areas of the *E. coli* genome

| Location | Albacore (default) |  | NanoReviser |  | NanoReviser<br>(methylation) |  |
| --- | --- | --- | --- | --- | --- | --- |
|  | Error<br>rate | Mismatch<br>rate | Error<br>rate | Mismatch<br>rate | Error<br>rate | Mismatch<br>rate |
| Overall | 19.55 | 7.74 | 14.22 | 4.69 | 12.05 | 3.29 |
| Methylated<br>bases | 34.69 | 18.54 | 29.53 | 24.55 | 26.27 | 21.38 |
| m6A bases | 37.32 | 32.97 | 32.45 | 28.41 | 28.38 | 24.89 |
| m4C bases | 31.89 | 25.66 | 27.08 | 23.60 | 25.72 | 22.53 |
| Modified bases | 33.46 | 26.27 | 28.13 | 22.43 | 25.12 | 19.25 |

The above results taken together indicated minimizing deletion errors to be at odds with minimizing insertion errors in training on low-coverage data or local data, and indicated methylation information to be crucial to the process of translating the electrical signal into a DNA sequence. Notably, decreasing the mismatch error was found to be the core improvement of NanoReviser (methylation), consistent with the high miscalling of the methylated bases.

### 3.5 Runtime comparison

We have compared the computing speeds on a local PC (MacBook Air, 2017) with one CPU processor (2.2 GHz Intel Core i7) and DDR3 memory (8GB, 1600 MHz), as well as on a high-performance computer (Dell R930, 2019) with four CPU processors (2.1 GHz Intel Xeon E7-4809 v4), four GPU processors (Nvidia Tesla T4) and 18 DDR4 memories (18×32G, 2133 MHz). The results are listed in Table 7. An advantage of Chiron and Scrappie in the tests was its use of parallel computation to accelerate the basecalling process, which was also used in NanoReviser basecalling process. Notably, since Scrappie needs to run on a Linux Ubuntu system, we just compared Scrappie raw and Scrappie events on the high-performance computer (Dell R930, 2019). With the local PC, running the tests using NanoReviser was over 40 times faster than that using Chiron, and much faster than DeepNano. When running the tests on the high-performance computer, NanoReviser was twice faster than Chiron. Due to the efficient operation ability of C, Scrappie, which was code in C, run faster than tools written in Python, including DeepNano, Chiron as well as NanoReviser. However, tools, which written in Python, have an excellent cross platform capability. Notably, some researchers (Rang et al., 2018) believed that the high demand of computing resources primarily limited the wide use of Chiron despite its high accuracy on basecalling. However, NanoReviser, which takes advantage of more of the available sequencing information than does Chiron, is a better solution for obtaining high-quality nanopore sequencing reads, especially when the computing resources are limited.

**Table 7.**
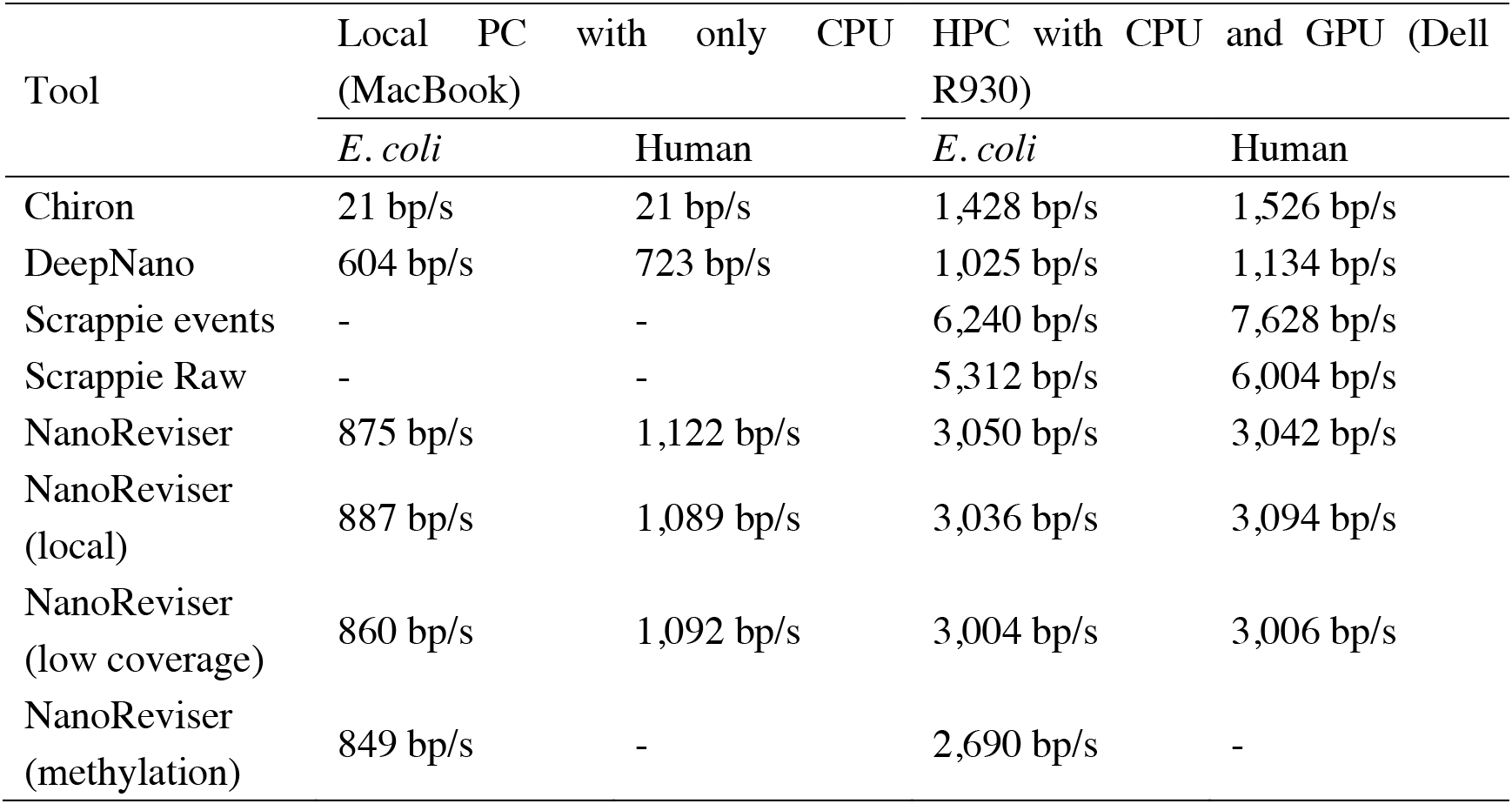
Running speeds of various tools used on E. coli data and human data

## 4 Conclusions

We have developed software based on a deep-learning algorithm and named NanoReviser, which was shown to efficiently correct the basecalling errors introduced by current basecallers provided by default. NanoReviser was validated using *E. coli* and *H. sapiens* data, and showed a higher accuracy than that of default basecaller, Albacore, the commonly regarded best basecaller, Chiron, as well as two separate models in Scrappie. Based on our elaborate training data construction, we found that NanoReviser learned more about insertion errors when trained on low-coverage data and more about deletion errors when trained on local reads. When trained on a combination of low-coverage data and local data, NanoReviser learned a compromise solution, which showed the importance of the training data selection. Moreover, due to our belief in the power of using genomics information when constructing a bioinformatics algorithm, we analyzed the methylated area of the *E. coli* genome. Here we found that the overall error rate for the modified area showed a different pattern than did the overall error rate for the whole genome. Specifically, the error rate was much higher for the modified areas, and mismatch error type contributed the most to the error rate. In order to handle the high mismatch error rate of the methylated bases, we used the methylation annotation to train NanoReviser. This training improved the performance of NanoReviser for both the methylated bases and on the whole genome scale. And when taking the methylation information into consideration, NanoReviser showed a reduced deletion error. Therefore, it may be concluded that there could be an enhanced relationship between modification detection, at least methylation detection, and basecalling, and we are looking forward to combining modification calling and basecalling. Furthermore, we believe that the error rate in the homopolymer region is a unique challenge in improving the basecalling accuracy, and according to the assessment on NA12878 chromosome 4, which contains the most homopolymer regions in all the NA12878 chromosomes, NanoReviser showed the ability to improve the basecalling accuracy on the homopolymer regions. Finally, we compared the running speeds of NanoReviser with DeepNano, Chiron and Scrappie on both our local PC and a high-performance computer. NanoReviser ran faster than did DeepNano and Chiron on the local PC and on the high-performance computer. On the high-performance computer, both Scrappie events and Scrappie raw are more efficient than NanoReviser. However, Scrappie can only supported by Linux Ubuntu system. Therefore, NanoReviser is expected to be a universal solution for the elimination of nanopore sequencing errors in the future.

## 5 Availability

NanoReviser is an open source program and the source code is available at https://github.com/pkubioinformatics/NanoReviser

## 7 Author Contributions

HQZ, LTW and LQ conceived the study; LTW developed the code; LTW, LSY and YYW performed data analysis under the supervision of HQZ; and LTW, LQ and HQZ wrote the manuscript. All authors read and approved the final manuscript.

## 9 Acknowledgments

The authors thank Dr. Binbin Lai, Dr. Qi Wang, Dr. Xiaoqi Wang, Dr. Yongchu Liu and Dr. Yang Li for their beneficial discussions and assistance with this paper.

